# Specialist–Generalist Trade-Offs in Microbial Growth Rates Across Soil Habitats

**DOI:** 10.1101/2025.05.20.655198

**Authors:** Megan M. Foley, Noah Sokol, Bram Stone, Steven J. Blazewicz, Katerina Estera-Molina, Alex Greenlon, Michaela Hayer, Jeffrey A. Kimbrel, Jennifer Pett-Ridge, Bruce A. Hungate

## Abstract

Microbial ecological strategies are shaped by a fundamental trade-off: is it better to specialize and thrive in a narrow niche or generalize and persist across diverse environments? In soils, this trade-off is particularly relevant in the rhizosphere and detritusphere, where microorganisms encounter distinct resource inputs from living and decaying roots. Using H ¹ O quantitative stable isotope probing (qSIP), we measured in situ bacterial and fungal growth rates in the rhizosphere, the root detritusphere, and in the combined presence of rhizosphere + root detritus to test whether specialists—microbes growing in a single habitat—grow faster than generalists that persist across multiple environments. Specialists grew consistently faster than generalists, suggesting a trade-off between the breadth of environmental conditions a microorganism can tolerate and its ability to grow quickly in a specific habitat. This cost to broad niche adaptation was apparent for bacteria, but growth rates of fungal saprotrophs varied little between specialists and generalists, reflecting how fundamental differences in life-history strategies between bacteria and fungi can shape microbial responses to resource availability and habitat heterogeneity. Net relatedness and nearest taxon indices indicated total bacterial communities were phylogenetically clustered while specialist and generalist communities were phylogenetically random, suggesting that functional traits, not lineage, best predict ecological strategy. In annual grassland soils, fast-growing specialists may dominate ecosystem processes when resources abound, and slow-growing generalists may sustain element transformations when conditions shift; understanding this interplay is key to predicting soil-carbon trajectories.

## 1. Introduction

Ecological strategies of soil microorganisms are shaped by fundamental trade-offs arising from cellular and ecological constraints. For soil microorganisms, this may include trading niche breadth for peak performance under specialized conditions. For example, maintaining the full enzymatic repertoire necessary for catabolizing the extraordinary diversity of plant-derived compounds in soils is metabolically taxing^1^. A single microorganism is unlikely to allocate enough proteome space to produce and regulate the diverse carbohydrate-active enzymes required to deconstruct cellulose, hemicellulose, lignin, and an array of root exudates without incurring substantial costs to growth or regulatory precision^2^. These constraints suggest that microbial niche breadth may be inherently limited by the energetic and spatial costs of proteome allocation, transcriptional control, and genome maintenance^3^. In contrast, specialist taxa may optimize for high efficiency on specific substrates and thus gaining a competitive edge in stable or substrate-rich niches^4,5^. Although consistent with ecological theory positing that generalists trade peak performance for versatility, empirical validations of such tradeoffs are elusive^6^.

In the habitats surrounding living and decaying plant roots, microorganisms experience dynamic resource availability and intense competition for nutrients. In rhizosphere soils, specialists may have traits such as the ability to move towards and adhere to root surfaces, allowing for the efficient utilization of root exudates^7^. In the detritusphere, microbial specialists may have filamentous growth forms or the ability to penetrate into areas they are decomposing, allowing for the full exploitation of resources present in the detritusphere ^8,9^. Generalists can likely exploit a wider spectrum of substrates and tolerate fluctuating environmental conditions but may do so at the expense of performance in any one habitat. Thus, the trade-off between specialization and generalism may be underpinned by functional traits, like metabolic versatility, that enable generalists to persist across diverse habitats but incur costs to fitness in a single environment.

The ways microbes establish dominance in the rhizosphere or detritusphere shape both short-term nutrient dynamics and the accumulation of soil organic carbon (SOC). Plant carbon can enter soil through rhizodeposition and root detritus, together feeding the largest pool of actively cycling organic carbon^10,11^. Microorganisms in these habitats play crucial roles in transforming these plant inputs into SOC through their growth and metabolic activities^12^. As microorganisms grow, they decompose organic matter and generate various organic compounds such as lipids, proteins, polysaccharides, nucleic acids, and metabolites. These microbial products can associate with minerals and persist in soils, thereby constituting a substantial fraction of slow-cycling SOC^13^. Because changes in resource availability in these habitats can flip which strategy dominates at a given point in time, linking SOC dynamics to ecological strategies offers a mechanistic, scalable alternative to taxonomy alone^14^.

To accurately capture the trade-offs between ecological strategies, it is essential to measure not just presence but active growth—since only a subset of microbial taxa are actively proliferating at any given time. Quantitative stable isotope probing (qSIP) using H ¹ O provides a direct measure of microbial growth, allowing us to infer how ecological strategies affect microbial success across different habitats without altering the carbon substrates they consume^15,16^. In this study, we aim to characterize specialist versus generalist taxa in different soil microbial habitats of a semi-arid Mediterranean grassland soil. In a 12-week greenhouse study, we used H ^18^O qSIP to assess taxon-specific growth rates in three key microbial habitats where microorganisms mediate the transformation of root-derived carbon to SOC – the rhizosphere, the detritusphere, and the combined rhizosphere + detritusphere. While the rhizosphere and detritusphere are often conceptualized in isolation, plant roots typically grow in the vicinity of decaying plant residues, resulting in a habitat that is influenced by both the rhizosphere and the detritusphere^14^. We categorized taxa by ecological strategy—based on their capacity to grow in specific habitats—and then tested whether these categories could reliably predict their performance in the rhizosphere, detritusphere, and the combined rhizosphere + detritusphere. We hypothesized that taxa that uniquely grow in one habitat (specialists) and taxa that grow in multiple habitats (generalists) have adaptations for distinct ecological niches and that there is a trade-off between the breadth of environmental conditions a microorganism can tolerate and its ability to grow quickly in a specific habitat.

## 2. Methods

### 2.1 Microcosm experiment

We conducted a 12-week greenhouse study to investigate dynamics of microbial specialists and generalists in different soil habitats. Planted and unplanted soil microcosms were used to generate a rhizosphere, detritusphere, and a combined rhizosphere + detritusphere of a California annual grassland soil. Soils were collected from the ‘A’ mineral horizon (0-10 cm) below a stand of California annual grasses dominated by *Avena barbata* at the University of California Hopland Research and Extension Center (39°00.106′N, 123°04.184′W). Hopland Research and Extension Center rests in the foothills of the Mayacamas Mountains and encompasses topographically rugged rangelands on the unceded territory of the Šóqowa people - the ancestral lands of the Hopland Band of Pomo Indians. The site is a California annual grassland ecosystem with a Mediterranean climate (MAT max/min = 23/7 C; MAP = 956 mm yr^−1^). *Avena* spp. is the dominant vegetation at the site and soil is a Typic Haploxeralfs (or a Luvisol) of the Witherall-Squawrock complex; the soil pH is ∼5.6, and contains 45% sand, 36% silt, and 19% clay^17^.

Roots were removed and soil was sieved (2-mm) and packed to field bulk density (1.21 g cm^−3^) in rectangular acrylic microcosms (11.5 cm × 2.9 cm × 25.5 cm). For rhizosphere microcosms, three germinated *Avena barbata* seedlings (collected from Hopland in spring 2018) were planted evenly apart in the soil. To generate root detritus for detritusphere microcosms, *A. barbata* was grown in a mixture of sand and Hopland soil for seven weeks. Roots were then harvested, repeatedly washed, dried at room temperature, and then cut into 1-5 mm fragments. In detritusphere microcosms, a 28-μm mesh ‘detrisuphere’ bag was buried in the microcosm center, which contained ∼65 g of soil mixed with the fragments of *A. barbata* root detritus (0.013 g root detritus dry g soil^−1^). For rhizosphere + detritusphere mesocosms, three germinated *Avena barbata* were planted in soil mesocosms containing a ‘detritusphere’ mesh bag.

All microcosms were incubated in growth chambers (56 cm × 56 cm × 76 cm) at the Environmental Plant Isotope Chamber facility at the Oxford Tract Greenhouse at University of California Berkeley^18,19^. Chambers were maintained with a 16-h light period per day (from 6 a.m. to 10 p.m.), a maximum daytime and nighttime air temperature of 27 °C and 24 °C, respectively, and a CO_2_ concentration between 400 and 450 ppm. Soil moisture was maintained at ∼16% gravimetric soil moisture (± 0.3 standard error) by weighing all microcosms twice weekly and adjusting soil moisture by mass.

### 2.2 Sample collection and isotope incubations

At the end of the 12-week experiment, soils from rhizosphere microcosms were collected by first gently shaking root systems to discard loosely attached soil. Soil still clinging to the roots after gentle shaking was then sampled and defined as rhizosphere soil. Soils from detritusphere microcosms were collected from inside the 28-μm mesh bag. Soils from rhizosphere + detritusphere microcosms were collected by isolating living the root system within the detritusphere mesh bag and collecting soils that remained clinging to roots after gentle shaking. Approximately 1 g dry weight of soil was immediately frozen on dry ice and frozen at −80 °C for amplicon sequencing.

Approximately 1.50 g dry weight of soil was transferred into sterile weigh boats for quantitative stable isotope probing (qSIP) incubations with isotopically enriched water; a separate set of soils were weighed out identically for incubations with natural abundance water. Soils were air dried to 9% gravimetric moisture in a laminar flow hood at room temperature for a few hours prior to qSIP incubations. Soils were transferred to sterile 30mL serum vials and approximately 0.5 milliliter of isotopically enriched water (98.15 at% ^18^O–H_2_O) or natural abundance water (as a control) was pipetted onto the soil slowly and evenly and then gently mixed with the pipette tip, raising soil moisture content to 29% gravimetric moisture. Vials were immediately sealed and incubated at room temperature in the dark for 48 hours. At the end of the incubation, soils were immediately frozen on dry ice and stored at −80 °C.

### 2.3 Quantitative stable isotope probing

DNA was extracted from soils using Qiagen DNeasy PowerSoil Pro Kits following the manufacturer’s instructions. Three replicate extractions were conducted for each sample and then replicate DNA extracts were combined. Samples were subjected to a cesium chloride density gradient formed by physical density separation via ultracentrifuge as previously described ^20,21^. 5 μg of DNA in 150 μL 1xTE was mixed with 1.00 mL gradient buffer, and 4.60 mL CsCl stock (1.885 g/mL) with a final average density of 1.730 g/mL. Samples were loaded into 5.2 mL ultracentrifuge tubes and spun at 20 °C for 108 h at 176,284 RCFavg in a Beckman Coulter Optima XE-90 ultracentrifuge using a VTi65.2 rotor.

Automated fractionation was performed using Lawrence Livermore National Laboratory’s high-throughput SIP pipeline^22^ Ultracentrifuge tube contents were separated into 19 fractions (∼200 μL each) using an Agilent Technologies 1260 isocratic pump delivering water at 0.25 mL min^−1^ through a 25G needle inserted through the top of the ultracentrifuge tube. Tubes were mounted in a Beckman Coulter fraction recovery system with a side port needle inserted through the bottom. The side port needle was routed to an Agilent 1260 Infinity fraction collector. Fractions were collected in 96-well deep well plates.

Fraction density was quantified using a Reichart AR200 digital refractometer fitted with a prism covering to facilitate measurement from 5 μL, as previously described^23^. Fractionated DNA was purified and concentrated using a Hamilton Microlab Star liquid handling system programmed to automate previously described glycogen/PEG precipitations^24^. Washed DNA pellets were suspended in 40 μL of 1xTE and the DNA concentration of each fraction was measured using a PicoGreen fluorescence assay.

### 2.4 16S rRNA gene qPCR and sequencing

DNA was amplified in triplicate 10 μL reactions using primers 515 F and 806 R ^25,26^. Each reaction contained 1 μL sample and 9 μL of Phusion Hot Start II High Fidelity master mix (Thermo Fisher Scientific) including 1.5 mM additional MgCl_2_. PCR conditions were 95 °C for 2 min followed by 20 cycles of 95 °C for 30 S, 64.5°C for 30 S, and 72 °C for 1 min. The assay was performed on a CFX 384 (Bio-Rad, Hercules, CA). Triplicate PCR products were pooled and diluted 10X and used as a template in a subsequent dual indexing reaction that used the same primers including the Illumina flowcell adaptor sequences and 8-nucleotide Golay barcodes.

Resulting amplicons were purified with AMPure XP magnetic beads (Beckman Coulter) and quantified with a PicoGreen assay on a BioTek Synergy HT plate reader. Samples were pooled at equivalent concentrations, purified with the AMPure XP beads, and quantified using the KAPA Sybr Fast qPCR kit (Kapa Biosciences). Libraries were sequenced on an Illumina MiSeq instrument at Northern Arizona University’s Genetics Core Facility using a 300-cycle v2 reagent kit.

### 2.5 18S rRNA gene qPCR and ITS2 sequencing

DNA was amplified in triplicate 10 μL reactions using primers FR1 and FF390 ^27^. Each reaction contained 1.25 μM of the primers FR1/FF390, 1× QuantiTect SYBR Green PCR Master Mix (Thermo Fisher Scientific) and 2.5 mM MgCl_2_. Standard curves were generated using genomic *Saccharomyces cereviciae* DNA (ATTC 201389d-5). PCR conditions were 95 °C for 15 min followed by 35 cycles of 95 °C for 15 s, 50 °C for 30 s and 70 °C for 1 min. The assay was performed on a CFX 384 (Bio-Rad, Hercules, CA). Sequencing library preparation was similar to constructing 16S rRNA gene sequencing libraries. The primers 5.8S-Fun and ITS4-Fun^28^ were used in the two PCR steps in 10uL reactions containing 0.8 μM of each primer, 0.01 U µL^−1^, Phusion HotStart II Polymerase (Thermo Fisher Scientific), 1X Phusion HF buffer (Thermo Fisher Scientific), 1.5 mM MgCl_2_, 6 % glycerol, and 200 µM dNTPs. PCR conditions were 95 °C for 2 min; 20 cycles of 95 °C for 30 s, 55 °C for 30 s, and 60 °C for 1 min. Libraries were sequenced on an Illumina MiSeq instrument at Northern Arizona University’s Genetics Core Facility.

### 2.6 Sequence processing and qSIP analysis

Paired-end reads were filtered to remove phiX and other contaminants with bbduk v38.56 ^29^. Fastq files were then trimmed to retain nucleotides 5–140 for the 16S F/R amplicons, and 5– 245/175 for the ITS F and R reads, respectively, filtered for quality (maxEE = 2, truncQ = 2) and used to generate amplicon sequence variants (ASVs) with DADA2 v1.20^30^. Chimeric sequences were determined and removed using removeBimeraDenovo from DADA2. ASV taxonomy was determined using the RDP 16S rRNA gene database (training set 18) using RDP classifier v2.11, keeping classifications with greater than 50% confidence^31^ and with the UNITE v8.3 database for ITS amplicons^32^. Alignments and phylogenetic trees were built using MAFFT v7.490 and FastTree v2.1.10^33,34^.

Excess atom fraction (EAF) ^18^O of DNA^35,36^ was quantified using the package “qSIP2”^37^ in R version 4.4.0^38^. qSIP analysis was restricted to taxa that occurred in at least 2 (of 3-4) natural abundance replicates and in at least 2 (of 12) density fractions. These criteria were chosen to reduce the likelihood of falsely interpreting spurious density shifts as growth. 90% confidence intervals for the change in density between control and enriched replicates, and corresponding values of EAF ^18^O, were computed on a per-replicate basis using bootstrap resampling (with replacement, 1,000 iterations) of natural abundance control replicates.

Values of EAF ^18^O that were negative or above the theoretical maximum enrichment of microbial DNA (EAF_max_) are physically impossible and were considered outliers if variation among technical replicates was high (defined here as SD > 0.15). EAF_max_ is computed as the product of the isotopic composition of soil water in each incubation (determined as a function of the amount of 97 atom % ^18^O water added and total soil water content) and the fraction of oxygen atoms in newly synthesized DNA that are derived from environmental water, which was set to 0.6 ^36^. A density correction was performed to account for slight differences in the preparation of the CsCl density gradient solution of each replicate^39^ and any remaining negative estimates of EAF ^18^O were corrected to zero. Estimates of EAF ^18^O above EAF_max_ likely reflecting rapid microbial growth and assimilation of ^18^O from additional sources like organic matter or prey biomass. These values (0.17% of observations) were censored to EAF_max_-0.02^40^. The relative growth rate of each taxon was estimated using the EAF ^18^O of a taxon’s DNA and the duration of the incubation (*t*) in days as:

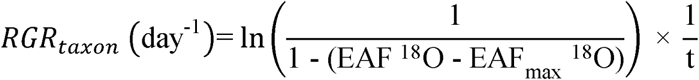

### 2.8 Statistical analysis

Amplicon sequencing of non-fractionated DNA demonstrated that qSIP quantified estimates of excess atom fraction (EAF) ^18^O for 92-94% of bacterial sequencing reads and 84-93% of fungal sequencing reads within each habitat (Supplementary Table 1 and Supplementary Figure 1). Venn diagrams and ternary plots were constructed and used to illustrate the distribution of growing ASVs, defined as taxa with positive relative growth estimates, across habitats. Ternary plots were produced using the ‘ggtern’ package in R ^41^.

To evaluate how microbial growth rates varied by habitat, ecological strategy, and their interaction, a two-way ANOVA was performed separately for bacterial and fungal taxa. Habitat (rhizosphere, detritusphere, combined) and ecological strategy (generalist vs. specialist) were included as fixed factors. Post hoc pairwise comparisons were conducted using t-tests.

Linear mixed-effects models were used to further evaluate the effects of habitat, ecological strategy, and taxonomic assignment on microbial growth rates. Residual diagnostics indicated an expected artifact from the applied censoring, resulting in a visible horizontal band and model false convergence. These data (.17% of observations of bacterial growth rates and .03% of observations of fungal growth rates) were treated as missing, and analyses were conducted on the remaining dataset. As a result, model slope and intercept estimates may underestimate effects at high growth rates. Models were fit using the nlme package (version 3.1-166) in R, with taxonomic assignment included as a nested random effect ( ∼1 | phylum/order/class/family/genus/ASV)^42^. To determine the optimal structure of the nested random term, a series of hierarchical linear mixed-effects models were constructed using restricted maximum likelihood (REML) estimation. All possible configurations of the nested random term were evaluated using Akaike Information Criterion corrected for finite sample size (AICc) and Bayesian Information Criterion (BIC). All models included habitat, ecological strategy, and their interaction as main effects. Once the optimal structure for the nested random term was identified, AICc and BIC were used to select the best statistical model from a set of candidate models that contained all possible combinations of the fixed effects while maintaining the selected nested random term structure. Finally, the mixed-effects model was compared against a simple linear model using AICc and BIC to quantify the effect of including taxonomy as a random term on model performance. Model selection was performed separately for bacterial vs. fungal growth rates. All models included a fixed variance structure to account for potential heteroscedasticity using the ‘varFixed’ function of the nlme package. ASVs with unknown taxonomic assignments, phyla represented by only one order, and families represented by fewer than three bacterial ASVs or fewer than two fungal ASVs were removed to reduce model overfitting ^43^. This limited our analysis to 862 bacterial and 156 fungal ASVs. Post hoc comparisons of least-squares means were conducted using the emmeans package to evaluate the main effects of habitat and ecological strategy^44^.

To evaluate the extent to which taxa growing in multiple habitats exhibited generalized growth responses to distinct plant carbon inputs (rhizodeposition versus root detritus), changes in growth rates were quantified for each taxon between the detritusphere and rhizosphere + detritusphere (rhizosphere effect on growth) and between the rhizosphere and rhizosphere + detritusphere (detritusphere effect on growth). Model II regression was used to assess how the effect of the rhizosphere on growth rates covaried with the effect of the detritusphere on growth rates of taxa growing in all habitats. Regressions were performed separately for fungal and bacterial ASVs using the ‘lmodel2’ package in R^45^. Because the rhizosphere effect and detritusphere effect share a common term (growth rate in the combined rhizosphere + detritusphere habitat), we benchmarked the observed slope and R² from the Model II regression against a permutation-based null. We permuted growth rates in the rhizosphere, detritusphere, and rhizosphere + detritusphere independently across taxa, recalculated rhizosphere and detritusphere effects, and recomputed the regression 9,999 times using the ‘lmodel2’ package in R. Upper- and lower-tail p-values were calculated to assess whether the observed slope and R² deviated significantly from the null expectation^46^.

The net relatedness index (NRI) and nearest taxon index (NTI) were calculated to evaluate phylogenetic clustering or overdispersion of microbial assemblages in the rhizosphere, detritusphere, and the combined rhizosphere-detritusphere. These indices were computed on a per-sample basis across multiple taxonomic groups: the entire set of taxa inferred from amplicon sequencing of unfractionated DNA, specialist taxa, generalist taxa. NRI and NTI were calculated using the picante package (version 1.8.2) in R^47^. All statistical tests were evaluated at an alpha level (α) of 0.10 and were conducted in R version 4.4.2^38^.

## 3. Results

### 3.1 Distribution of growing microbial taxa across soil habitats

The majority of bacterial amplicon sequence variants (ASVs) grew in multiple habitats (54%; Figure 1 and Supplementary Figure 1) or exclusively in the rhizosphere (40%). A smaller proportion of bacterial ASVs grew exclusively in the detritusphere (2%) or the combined rhizosphere + detritusphere (4%). In contrast, most fungal ASVs were detected growing in multiple habitats (66%; Figure 1) with fewer ASVs restricted to the rhizosphere (15%), the detritusphere (4%), or the combined rhizosphere + detritusphere (15%).

**Figure 1:**
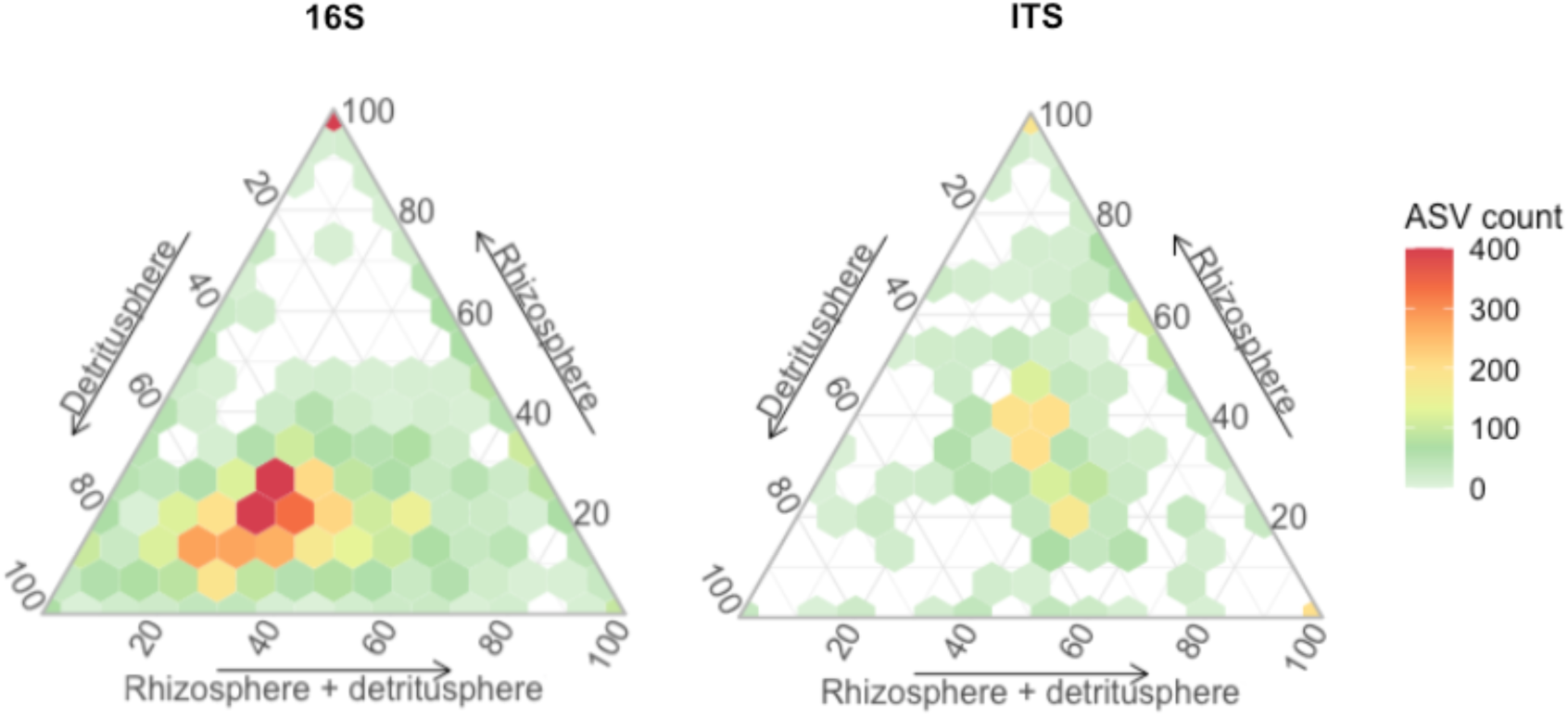
Distribution of relative growth rates for amplicon sequence variants (ASVs) across three habitats: rhizosphere soil, detritusphere soil, and the combined rhizosphere + detritusphere. Each hexagon represents a compositional bin and color indicates the number of ASVs whose growth rate proportions fall within that bin. Axes are scaled from 0% to 100%, so a vertex denotes exclusive growth in one habitat, whereas the center (33 %, 33 %, 33 %) indicates equal growth in all three. ASVs situated at a vertex highlight habitat specialists (growth detected in a single habitat).

**Figure 2:**
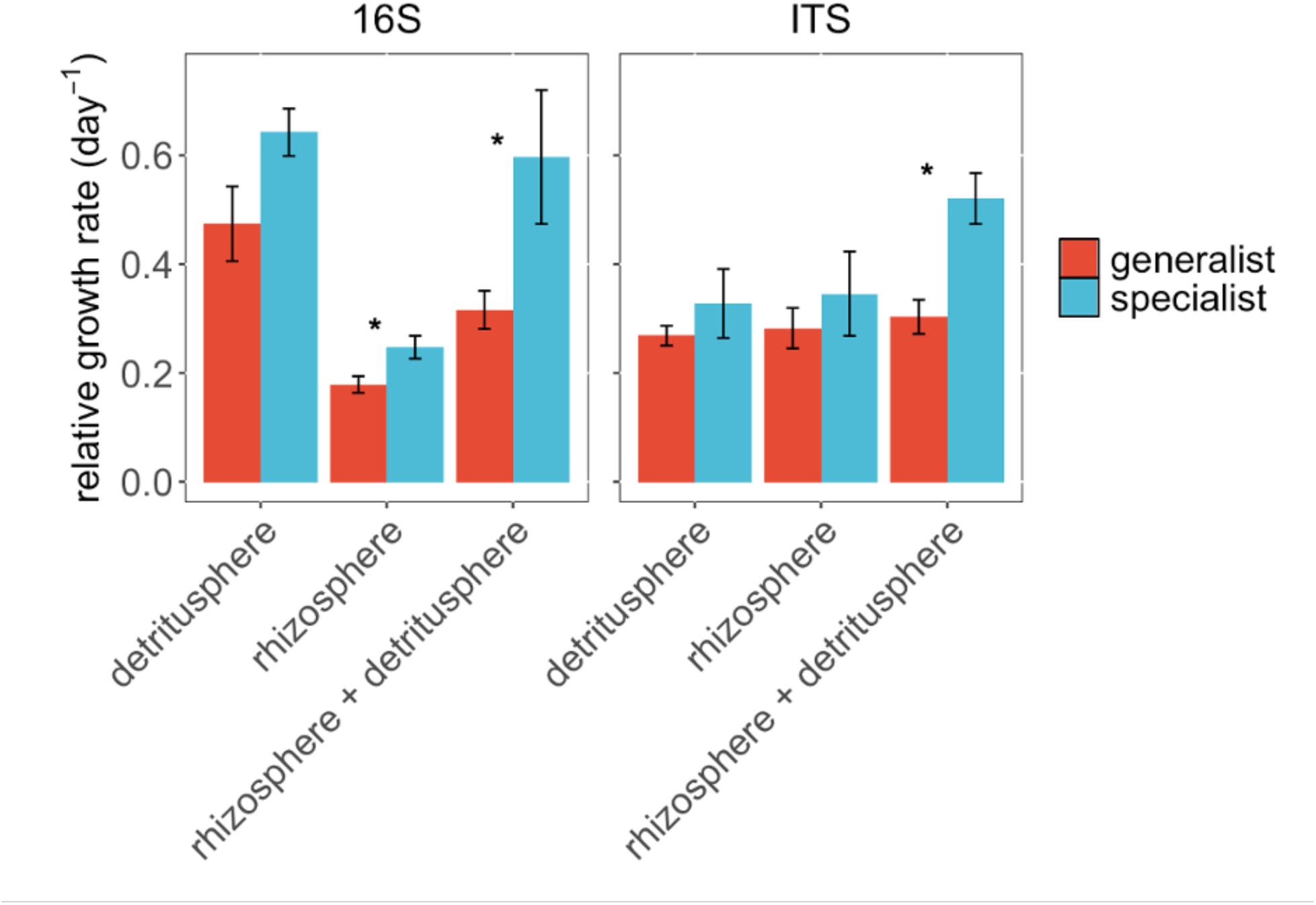
Relative growth rates of microbial generalists and specialists across habitats. Specialists were defined as taxa uniquely growing in one habitat; generalists grew in multiple habitats and were assayed in each. Two-way ANOVAs tested for effects of habitat and ecological strategy on growth rates, separately for bacteria and fungi. For bacteria, growth rates differed by strategy (*F* , = 10.8, *p* = 0.005) and habitat (*F* , = 14.9, *p* < 0.001); specialists grew faster in the rhizosphere (t-test, *p* = 0.04) and the combined habitat (t-test, *p* = 0.10). Fungal growth also differed by strategy (*F* , = 8.06, *p* = 0.01), with higher specialist growth in the combined habitat (t-test, *p* = 0.01). Bars show standard error; asterisks indicate *p* < 0.10 (t-tests).

### 3.2 Bacterial growth rates

We classified ASVs into two ecological strategies: “generalists,” which grow across multiple habitats, and “specialists,” which are restricted to a single habitat. Bacterial growth rates differed by ecological strategy (two-way ANOVA: *F*_1,16_ = 10.8, *p* = 0.005; Figure 3) and habitat (two-way ANOVA: *F*_2,16_ = 14.9, *p* < 0.001; Figure 3). Specialists grew faster than generalists in the rhizosphere (t-test, *p* = 0.04) and in the combined rhizosphere + detritusphere (t-test, *p* = 0.10) and showed non-significant trends toward faster growth in the detritusphere (t-test, *p* = 0.12). On average, rhizosphere specialists grew slower than detritusphere (t-test*, p* = 0.004) and rhizosphere + detritusphere (t-test, *p* = 0.06) specialists. Similarly, rhizosphere generalists grew slower than detritusphere (t-test*, p* = 0.04) and rhizosphere + detritusphere (t-test, *p* = 0.02) generalists.

**Figure 3:**
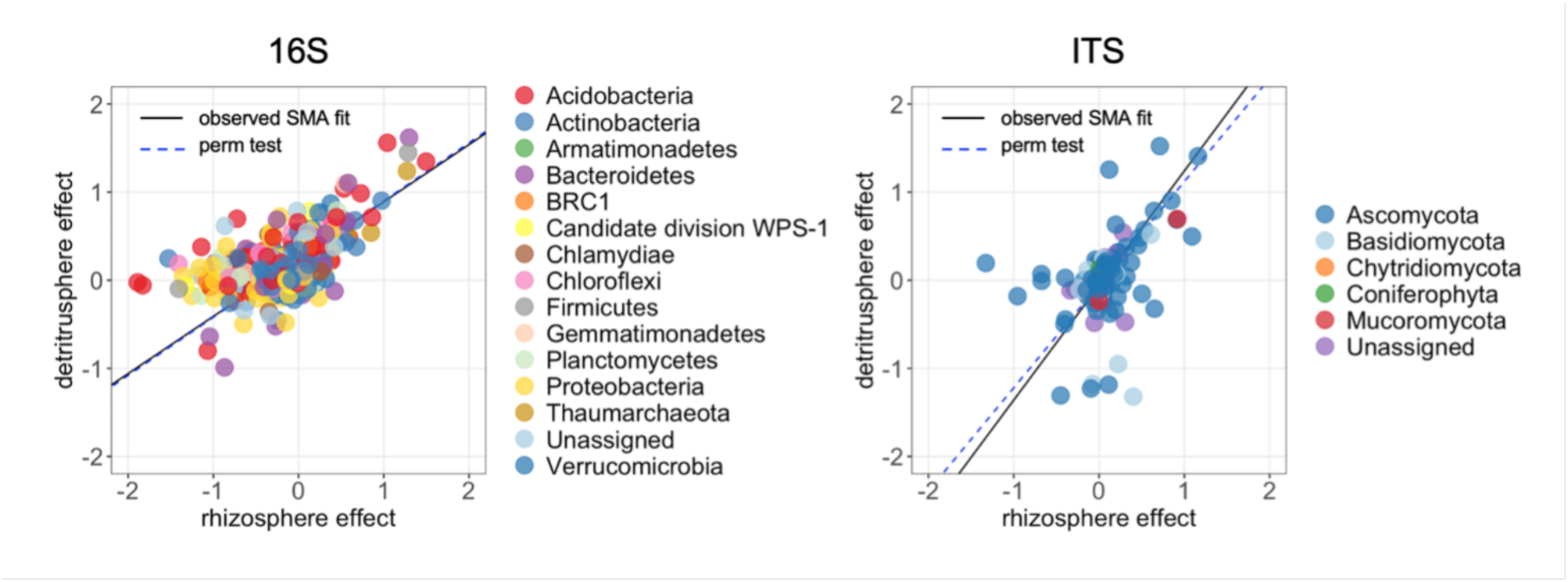
Relationship between the rhizosphere effect and detritusphere effect on taxon-specific growth rates. Each point represents an ASV detected in all three habitats; colors indicate phylum-level taxonomy. The solid line shows the observed standard major axis (SMA) regression fit (type II), and the dashed line shows the mean SMA fit across 9,999 permutations in which growth rates in each habitat were independently shuffled across ASVs. For fungi, the observed slope exceeded the permutation expectation (P = 0.039), indicating a weak but consistent generalist response. For bacteria, the observed slope and R² did not differ from the permutation-based null (P > 0.4 and P = 0.53, respectively), suggesting that any apparent coupling between the rhizosphere effect and the detritusphere effect was driven by the shared-term artifact.

Linear mixed-effects modeling confirmed the importance of habitat in explaining bacterial growth rates (*F* , = 227.56, *p* < 0.001), while ecological strategy alone did not significantly improve the model fit (*F* , = 0.42, *p* = 0.52). However, pairwise comparisons within the model indicated that specialists exhibited higher growth rates than generalists (*t* = 4.37, *p* < 0.001). The inclusion of taxonomic structure as a nested random effect (phylum, order, and ASV; see Supplementary Results) significantly improved model performance (ΔAICc = 1536.74; ΔBIC = 1517.22).

### 3.3 Fungal growth rates

Fungal growth rates differed by ecological strategy (two-way ANOVA: *F*_1,16_ = 8.06, *p* = .01; Figure 3) and habitat (two-way ANOVA: *F*_2,16_ = 2.97, *p* = 0.08; Figure 3). Post-hoc testing indicated specialists exhibited higher growth rates than generalists in the combined rhizosphere + detritusphere habitat only (t-test, *p* = 0.01). Mixed-effects modeling supported the significance of both habitat (*F* , = 7.90, *p* < 0.001) and ecological strategy (*F* , = 4.50, *p* = 0.04) in explaining fungal growth rates. The inclusion of taxonomic structure as a random effect significantly improved model performance (ΔAICc = 356.77; ΔBIC = 352.01), though the explained variance was relatively low (conditional *R*² = 0.05).

### 3.2 Consistency of Growth Responses to Rhizodeposition vs. Root Detritus

To test whether taxa present in all three habitats exhibit generalized growth responses to rhizodeposition and root detritus, we calculated two habitat effects for each ASV: the rhizosphere effect (RE = growth rate_rhizosphere + detritusphere_ – growth rate_detritusphere_) and the detritusphere effect (DE = growth rate_rhizosphere + detritusphere_ – growth rate_rhizosphere_). We used linear regression to assess how the effect of the rhizosphere on growth rates covaried with the effect of the detritusphere on growth rates of taxa growing in all habitats. For bacteria, the observed RE– DE slope (β_obs_ = 0.28 ± 0.02 SE) was indistinguishable from a permutation mean (β_null_ = 0.29, P>0.4). R² also did not exceed the null expectation (*p* = 0.53). Any apparent coupling can thus be fully explained by the shared-term artifact. For fungi, the observed slope exceeded the permutation expectation (β_obs_ = 0.47 vs β_null_ = 0.33; *p*_upper_ = 0.039). The overall fit (R²_obs_ = 0.11) was no tighter than random (*p* = 0.90), suggesting substantial taxon-to-taxon variation around this trend.

### 3.4 Phylogenetic structure of specialist and generalist communities

Net relatedness index (NRI) and nearest taxon index (NTI) were used to assess the phylogenetic structure of microbial communities in each habitat. Net relatedness index (NRI) and nearest taxon index (NTI) did not differ from zero for specialist microbes in any habitat, indicating that this subset of the total community was phylogenetically random. The same was also true for generalist microbes (Figure 4). Total bacterial communities—defined as the collection of ASVs obtained from sequencing unfractionated DNA—exhibited pronounced phylogenetic clustering, as reflected by positive NRI and NTI values. For total fungal communities, NRI values remained at zero, while NTI values were moderately positive, suggesting slight phylogenetic clustering, particularly at finer taxonomic scales.

**Figure 4:**
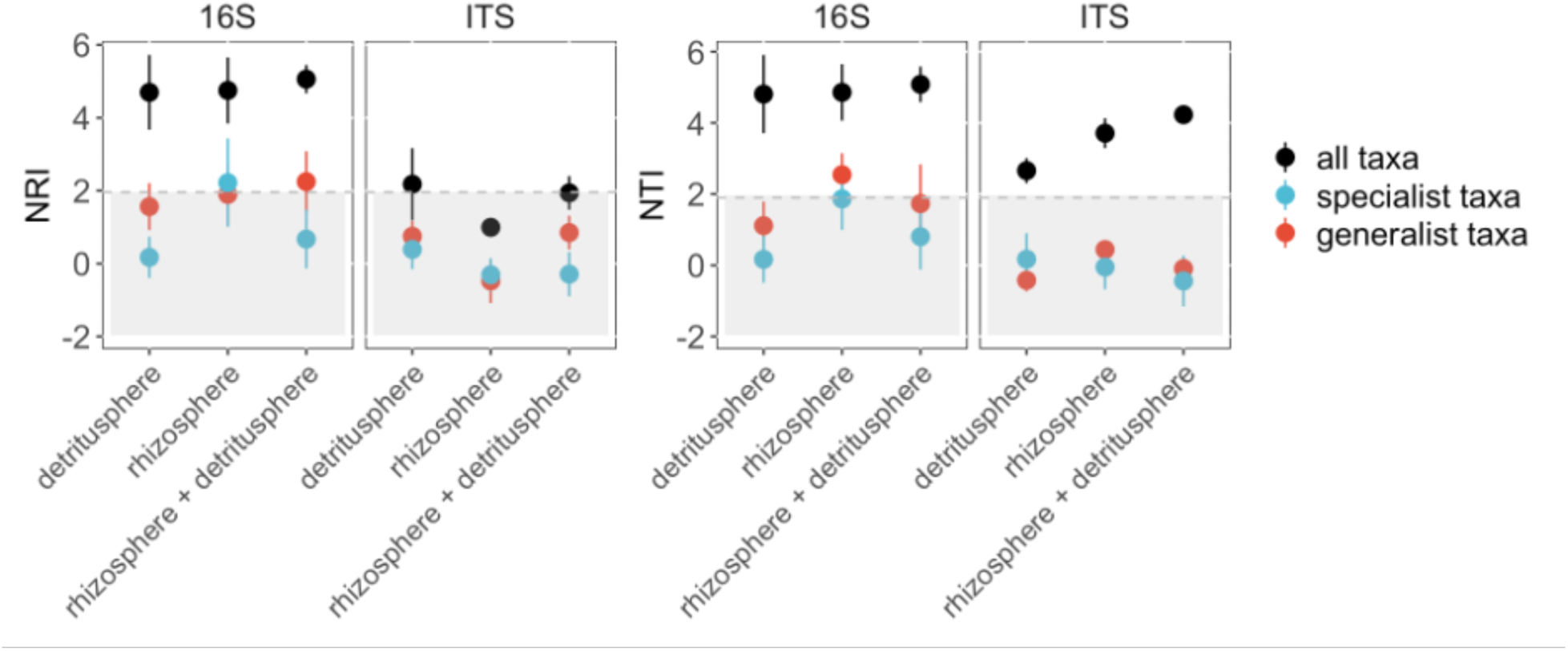
Net relatedness index (NRI) and nearest taxon index (NTI) of total, specialist, and generalist microbial communities across habitats. Total communities were inferred from 16S and ITS amplicon sequencing of unfractionated DNA (black points). Error bars indicate standard error. Points outside the grey shaded region differ significantly from random expectations (95% confidence). Total bacterial communities showed strong phylogenetic clustering (positive NRI and NTI), while fungal communities exhibited weak clustering (NTI only). Specialist and generalist taxa did not deviate from random phylogenetic structure.

## 4. Discussion

Microbial growth rates fundamentally shape element cycling in soil, yet we lack in-situ measurements that tie growth to ecological strategies. We used quantitative stable isotope probing to assess soil microbial growth in the three dominant belowground habitats where microorganisms mediate the transformation of root-derived carbon to soil organic carbon: the rhizosphere, the detritusphere, and the combined rhizosphere + detritusphere. Our results demonstrate that soil bacteria experience a fundamental trade-off between generalism and specialization. Specialists achieved higher growth rates than generalists in their respective single-habitat environments, indicating the fitness gains of a narrower niche (Figure 3).

Our data are consistent with broader ecological theory positing that organisms with wide niche breadth forgo peak performance in any single setting to gain adaptive flexibility across many^48–50^. Organisms adapted to diverse environments often evolve metabolic flexibility^4^, allowing them to exploit a wide range of resources, or rely on complex regulatory networks to fine-tune gene expression in response to changing conditions^51^. The metabolic versatility of generalists may come at the cost of reduced efficiency in utilizing resources abundant in a particular habitat or a slower response to signals unique to the rhizosphere or detritusphere^4^.

Specialists, by contrast, may prioritize traits that optimize fast growth within a specific niche^52^ Such specialization is well documented in the rhizosphere, where resource competition and the unique composition of root exudates often select for microbial taxa with specific metabolic pathways or rapid chemotactic responses, and the detritusphere, which favors microorganisms with specialized enzymatic repertoires for lignocellulose decomposition^53–55^. Notably, the advantage of specialists in the combined rhizosphere and detritusphere habitat suggests they benefit from resource complementarity or synergy^56^.

In our study, differences in growth rates between specialists and generalists were more pronounced for bacteria compared to fungal saprotrophs. Fungal generalists did not show a strong disadvantage in growth rate relative to fungal specialists (Figure 3), suggesting that saprotrophic fungi may be less affected by the trade-off that constrains bacteria. These findings are consistent with broader ecological patterns suggesting that bacteria often exhibit rapid, resource-driven responses to environmental change—particularly in the rhizosphere where exudates can stimulate growth^57–59^. Fungi, in contrast, through their filamentous networks and wide array of hydrolytic and oxidative enzymes, can access and decompose a variety of substrates over broader spatial scales and timeframes^8,55,60^. Fungal saprotrophs may therefore exploit diverse resources without substantially compromising growth rate in a specific habitat^43,61,62^. The idea that hyphal foraging abilities and broad enzyme repertoires reduce the fitness cost of niche breadth is supported by our finding that fungal generalists gain modestly from both rhizodeposits and root litter sources (Figure 1). In contrast, the RE–DE relationship for bacteria was statistically indistinguishable from the permutation null, leaving open whether bacterial generalists truly face a strong trade-off or whether our metric was simply confounded by the shared RD term. Adding a bulk-soil control (growth in neither roots nor detritus) in a fully factorial design would allow us to compute independent response ratios and eliminate the coupling problem altogether.

Our phylogenetic analyses provide further insight into the nature of specialist and generalist strategies. We observed that total bacterial communities in each habitat were phylogenetically clustered, but subsets of specialists and generalists themselves were phylogenetically random. In other words, closely related taxa did not necessarily share the same strategy. This result suggests that taxonomic identity helps predict which taxa initially recruit to these belowground habitats, but that whether a taxon behaves as a specialist or generalist is independent of phylogeny and may depend more on functional traits^63^. The decoupling of phylogeny and function in this context underscores the idea that diverse lineages can end up with similar ecological strategies, reinforcing the value of trait-centric approaches in microbial ecology^64,65^.

From a biogeochemical perspective, our findings highlight how trade-offs might scale up to influence soil carbon cycling under changing resource conditions. When inputs of root exudates and plant litter fluctuate, the balance of generalist and specialist microbes could govern the rate and continuity of decomposition. A community enriched in generalists may sustain more continuous decomposition across variable conditions while communities dominated by specialists may rapidly metabolize substrates only when resources become abundant. Moving forward, it will be critical to quantify microbial growth rates across habitats independent of immediate carbon availability, given that microbial growth, turnover, and carbon use efficiency are theorized to be key determinants of soil organic carbon formation^9,66^. However, the current study has several caveats. The relatively rapid growth observed in the detritusphere may partly reflect rewetting effects during qSIP incubations, such as the mobilization of dissolved organic carbon following water additions^67^. Similarly, the rhizosphere growth rates measured at the end of the growing season likely reflect a period of lower microbial activity due to reduced rhizodeposition. Future work spanning different points in plant phenology and refining moisture controls should allow future studies to resolve these dynamics with greater precision.

Our results advance our understanding of microbial niche breadth and trait-based ecology by illustrating tradeoffs in ecological strategies that may influence how microbial communities respond to resource variability. In summary, bacterial generalists demonstrated wider niche breadth at the expense of rapid growth compared to bacterial specialists, underscoring context-dependent advantages of broad versus narrow niche strategies. Fungal generalists exhibited minimal evidence of such trade-offs, potentially due to enzymatic and structural traits that extend their functional niche. Notably, specialist and generalist growth strategies were phylogenetically random, indicating convergent life-history adaptations across lineages. In soils with changing resources, slower growing generalists may sustain element transformations when conditions shift while faster growing specialists dominate during periods of resource abundance. Understanding this interplay is key to predicting soil-carbon trajectories. Accounting for such strategies may improve predictions of soil carbon trajectories in the face of changing plant inputs.

## Acknowledgements

We appreciate discussion with members of the Lawrence Livermore National Laboratory (LLNL) Soil Microbiome Scientific Focus Area team. This work was supported by the U.S. Department of Energy’s (DOE) Biological Systems Science Division Program in Genomic Science (LLNL ‘Microbes Persist’ Soil Microbiome Scientific Focus Area SCW1632). Work at LLNL was conducted under the auspices of the US Department of Energy under Contract DE-AC52-07NA27344.

