## Supplementary Information for "Specialist–Generalist Trade-Offs in Microbial Growth Rates Across Soil Habitats"

**Supplementary Methods**

Linear mixed effects model analysis

We compared a linear model without taxonomic information against a mixed-effects model with taxonomy coded as a nested random effect to assess the extent to which taxonomy influences growth rates. Model performance improved significantly with the inclusion of taxonomy as a nested random effect in the mixed-effects model for both bacterial (ΔAICc = 1536.74; ΔBIC = 1517.22) and fungal growth rates (ΔAICc = 356.77; ΔBIC = 352.01). For bacterial growth rates, the optimal nested random effect structure included phylum, order, and ASV (Supplementary Table 2) and the best statistical model included habitat and ecological strategy as fixed effects (Supplementary Table 3; conditional *R*^2^ = .54, habitat: *F*_2, 4074_ = 227.56, *p* < .001, ecological strategy: *F*_1, 807_ = 0.42, *p* = .52). For fungal growth, the optimal random effect structure included ASV (Supplementary Table 4) and the best statistical model included habitat and ecological strategy as fixed effects (Supplementary Table 5; conditional *R*^2^ = .05, habitat: *F*_2,681_ = 7.90, *p* < .001, ecological strategy: *F*_1,154_ = 4.50, *p* = .04).

**Supplementary Tables**

| Habitat | Marker Gene | Proportion of reads with resolvable estimates of EAF ^18^O |
| --- | --- | --- |
| Rhizosphere | 16S | 94 ± 0.5% |
| Rhizosphere | ITS | 84 ± 5% |
| Detritusphere | 16S | 92 ± 2% |
| Detritusphere | ITS | 93 ± 4% |
| Rhizosphere + detritusphere | 16S | 93 ± 1% |
| Rhizosphere + detritusphere | ITS | 85 ± 3% |

**Supplementary Table 1**: Proportion (average ± standard deviation) of amplicon sequencing reads with resolvable estimtes of EAF ^18^O for bacterial (16S) and fungal (ITS) taxa in each soil habitat.

| **Random term** | ***df*** | **ΔAICc** | **ΔBIC** |
| --- | --- | --- | --- |
| phylum/class/order/family/genus/ASV | 13 | 6.0 | 25.5 |
| phylum/class/order/family/genus | 12 | 534.2 | 547.2 |
| phylum/class/order/family/ASV | 12 | 4.0 | 17.0 |
| phylum/class/order/genus/ASV | 12 | 4.0 | 17.0 |
| phylum/class/family/genus/ASV | 12 | 17.9 | 30.9 |
| phylum/order/family/genus/ASV | 12 | 4.0 | 17.0 |
| class/order/family/genus/ASV | 12 | 8.8 | 21.8 |
| phylum/class/order/family | 11 | 605.7 | 612.2 |
| phylum/class/order/genus | 11 | 532.2 | 538.7 |
| phylum/class/family/genus | 11 | 538.5 | 545.0 |
| phylum/order/family/genus | 11 | 532.2 | 538.7 |
| class/order/family/genus | 11 | 536.3 | 542.8 |
| phylum/class/order/ASV | 11 | 2.0 | 8.5 |
| phylum/class/family/ASV | 11 | 15.9 | 22.4 |
| phylum/order/family/ASV | 11 | 2.0 | 8.5 |
| class/order/family/ASV | 11 | 6.8 | 13.3 |
| phylum/class/genus/ASV | 11 | 23.9 | 30.4 |
| phylum/order/genus/ASV | 11 | 2.0 | 8.5 |
| class/order/genus/ASV | 11 | 6.8 | 13.3 |
| phylum/family/genus/ASV | 11 | 16.0 | 22.6 |
| class/family/genus/ASV | 11 | 18.8 | 25.3 |
| order/family/genus/ASV | 11 | 7.6 | 14.2 |
| phylum/class/order | 10 | 706.5 | 706.5 |
| phylum/class/family | 10 | 609.2 | 609.2 |
| phylum/class/genus | 10 | 538.0 | 538.0 |
| phylum/order/family | 10 | 603.7 | 603.7 |
| phylum/order/genus | 10 | 530.2 | 530.2 |
| class/order/family | 10 | 607.3 | 607.3 |
| class/order/genus | 10 | 534.3 | 534.3 |
| phylum/family/genus | 10 | 537.4 | 537.4 |
| class/family/genus | 10 | 538.8 | 538.8 |
| order/family/genus | 10 | 535.1 | 535.1 |
| phylum/class/ASV | 10 | 57.7 | 57.7 |
| phylum/order/ASV | 10 | 0.0 | 0.0 |
| class/order/ASV | 10 | 4.8 | 4.8 |
| phylum/family/ASV | 10 | 14.1 | 14.1 |
| class/family/ASV | 10 | 16.8 | 16.8 |
| order/family/ASV | 10 | 5.6 | 5.6 |
| phylum/genus/ASV | 10 | 24.7 | 24.7 |
| class/genus/ASV | 10 | 23.8 | 23.8 |
| order/genus/ASV | 10 | 5.6 | 5.6 |
| family/genus/ASV | 10 | 21.9 | 21.9 |
| phylum/class | 9 | 1116.7 | 1110.2 |
| phylum/order | 9 | 704.5 | 698.0 |
| phylum/family | 9 | 607.2 | 600.7 |
| phylum/genus | 9 | 540.2 | 533.7 |
| class/order | 9 | 706.5 | 699.9 |
| class/family | 9 | 610.6 | 604.1 |
| class/genus | 9 | 537.8 | 531.3 |
| order/family | 9 | 605.4 | 598.9 |
| order/genus | 9 | 533.1 | 526.6 |
| family/genus | 9 | 543.6 | 537.1 |
| phylum/ASV | 9 | 62.2 | 55.7 |
| class/ASV | 9 | 57.9 | 51.4 |
| order/ASV | 9 | 3.6 | -2.9 |
| family/ASV | 9 | 19.9 | 13.4 |
| genus/ASV | 9 | 35.6 | 29.1 |
| phylum | 8 | 1333.3 | 1320.3 |
| class | 8 | 1114.7 | 1101.7 |
| order | 8 | 704.5 | 691.4 |
| family | 8 | 611.9 | 598.9 |
| genus | 8 | 550.0 | 537.0 |
| ASV | 8 | 100.7 | 87.7 |

**Supplementary Table 2:** We used AICc and BIC criteria to evaluate all possible configuarations of a nested random term for hierarchical taxonomic assignments of bacterial taxa: phylum, class, order, family, genus, amplicon sequence variant (ASV). In all models, habitat and ecological strategy, and their interaction, were included as fixed effects. **Δ**AICc and **Δ**BIC values indicate the change in these criteria relative to the optimal nested random effect structure (u_phylum/order/ASV_ ).

| **Main effects** | ***df*** | **ΔAICc** | **ΔBIC** |
| --- | --- | --- | --- |
| Habitat:Ecological strategy, Habitat, Ecological strategy | 10 | 2.9 | 15.9 |
| Habitat, Ecological strategy | 8 | 0.0 | 0.0 |
| Habitat | 7 | 17.0 | 10.5 |
| Ecological strategy | 6 | 425.1 | 412.1 |

**Supplementary Table 3:** We progressively dropped main effect terms from model predicting bacterial growth rates and used AICc and BIC criteria to evaluate model parsimony. **Δ**AICc and **Δ**BIC values indicate the change in these criteria relative to the most parsimonious model.

| **Random term** | ***df*** | **ΔAICc** | **ΔBIC** |
| --- | --- | --- | --- |
| phylum/class/order/family/genus/ASV | 13 | 8.3 | 32.0 |
| phylum/class/order/family/genus | 12 | 52.3 | 71.2 |
| phylum/class/order/family/ASV | 12 | 6.3 | 25.2 |
| phylum/class/order/genus/ASV | 12 | 6.3 | 25.2 |
| phylum/class/family/genus/ASV | 12 | 7.2 | 26.1 |
| phylum/order/family/genus/ASV | 12 | 6.3 | 25.2 |
| class/order/family/genus/ASV | 12 | 6.3 | 25.2 |
| phylum/class/order/family | 11 | 62.5 | 76.6 |
| phylum/class/order/genus | 11 | 50.3 | 64.5 |
| phylum/class/family/genus | 11 | 51.8 | 66.0 |
| phylum/order/family/genus | 11 | 50.3 | 64.5 |
| class/order/family/genus | 11 | 50.3 | 64.5 |
| phylum/class/order/ASV | 11 | 4.3 | 18.5 |
| phylum/class/family/ASV | 11 | 5.2 | 19.4 |
| phylum/order/family/ASV | 11 | 4.3 | 18.5 |
| class/order/family/ASV | 11 | 4.3 | 18.5 |
| phylum/class/genus/ASV | 11 | 5.8 | 19.9 |
| phylum/order/genus/ASV | 11 | 4.3 | 18.5 |
| class/order/genus/ASV | 11 | 4.3 | 18.5 |
| phylum/family/genus/ASV | 11 | 5.2 | 19.4 |
| class/family/genus/ASV | 11 | 5.2 | 19.4 |
| order/family/genus/ASV | 11 | 4.3 | 18.5 |
| phylum/class/order | 10 | 72.8 | 82.3 |
| phylum/class/family | 10 | 61.2 | 70.6 |
| phylum/class/genus | 10 | 50.7 | 60.1 |
| phylum/order/family | 10 | 60.5 | 69.9 |
| phylum/order/genus | 10 | 48.3 | 57.8 |
| class/order/family | 10 | 60.5 | 69.9 |
| class/order/genus | 10 | 48.3 | 57.8 |
| phylum/family/genus | 10 | 49.8 | 59.3 |
| class/family/genus | 10 | 49.8 | 59.3 |
| order/family/genus | 10 | 48.3 | 57.8 |
| phylum/class/ASV | 10 | 4.0 | 13.5 |
| phylum/order/ASV | 10 | 2.3 | 11.8 |
| class/order/ASV | 10 | 2.3 | 11.8 |
| phylum/family/ASV | 10 | 3.2 | 12.6 |
| class/family/ASV | 10 | 3.2 | 12.6 |
| order/family/ASV | 10 | 2.3 | 11.8 |
| phylum/genus/ASV | 10 | 3.8 | 13.2 |
| class/genus/ASV | 10 | 3.8 | 13.2 |
| order/genus/ASV | 10 | 2.3 | 11.8 |
| family/genus/ASV | 10 | 3.2 | 12.6 |
| phylum/class | 9 | 178.9 | 183.6 |
| phylum/order | 9 | 70.8 | 75.6 |
| phylum/family | 9 | 59.2 | 63.9 |
| phylum/genus | 9 | 48.7 | 53.4 |
| class/order | 9 | 70.8 | 75.6 |
| class/family | 9 | 59.2 | 63.9 |
| class/genus | 9 | 48.7 | 53.4 |
| order/family | 9 | 58.5 | 63.2 |
| order/genus | 9 | 46.3 | 51.1 |
| family/genus | 9 | 47.8 | 52.5 |
| phylum/ASV | 9 | 2.0 | 6.7 |
| class/ASV | 9 | 2.0 | 6.7 |
| order/ASV | 9 | 0.3 | 5.1 |
| family/ASV | 9 | 1.2 | 5.9 |
| genus/ASV | 9 | 1.8 | 6.5 |
| phylum | 8 | 330.8 | 330.8 |
| class | 8 | 177.4 | 177.4 |
| order | 8 | 68.8 | 68.8 |
| family | 8 | 57.2 | 57.2 |
| genus | 8 | 46.7 | 46.7 |
| ASV | 8 | 0.0 | 0.0 |

**Supplementary Table 4:** We used AICc and BIC criteria to evaluate all possible configuarations of a nested random term for hierarchical taxonomic assignments of fungal taxa: phylum, class, order, family, genus, amplicon sequence variant (ASV). In all models, habitat and ecological strategy, and their interaction, were included as fixed effects. **Δ**AICc and **Δ**BIC values indicate the change in these criteria relative to the optimal nested random effect structure (u_ASV_ ).

| **Main effects** | ***df*** | **ΔAICc** | **ΔBIC** |
| --- | --- | --- | --- |
| Habitat:Ecological strategy, Habitat, Ecological strategy | 8 | 1.2 | 12.6 |
| Habitat, Ecological strategy | 6 | 0.0 | 2.0 |
| Habitat | 5 | 2.8 |  |
| Ecological strategy | 4 | 11.7 | 4.2 |

**Supplementary Table 5:** We progressively dropped main effect terms from model predicting fungal growth rates and used AICc and BIC criteria to evaluate model parsimony. **Δ**AICc and **Δ**BIC values indicate the change in these criteria relative to the optimal model.

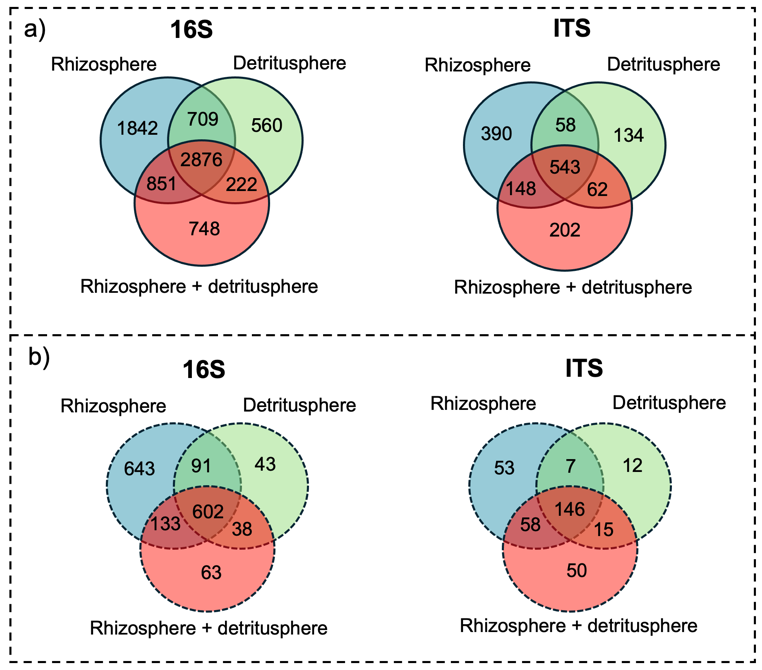

**Supplementary Figure 1:** Venn diagrams of the number of amplicon sequence variants (a) detected by 16S and ITS sequencing and (b) detected as growing via H218O quantitative stable isotope probing.
